# Production and secretion of functional full-length SARS-CoV-2 spike protein in *Chlamydomonas reinhardtii*

**DOI:** 10.1101/2021.12.13.472433

**Authors:** Anna Kiefer, Justus Niemeyer, Anna Probst, Gerhard Erkel, Michael Schroda

## Abstract

The spike protein is the major protein on the surface of coronaviruses. It is therefore the prominent target of neutralizing antibodies and consequently the antigen of all currently admitted vaccines against SARS-CoV-2. Since it is a 1273-amino acids glycoprotein with 22 N-linked glycans, the production of functional, full-length spike protein was limited to mammalian and insect cells, requiring complex culture media. Here we report the production of full-length SARS-CoV-2 spike protein – lacking the C-terminal membrane anchor – as a secreted protein in the prefusion-stabilized conformation in the unicellular green alga *Chlamydomonas reinhardtii*. We show that the spike protein is efficiently cleaved at the furin cleavage site during synthesis in the alga and that cleavage is abolished upon mutation of the multi-basic cleavage site. We could enrich the spike protein from culture medium by ammonium sulfate precipitation and demonstrate its functionality based on its interaction with recombinant ACE2 and ACE2 expressed on human 293T cells. *Chlamydomonas reinhardtii* is a GRAS organism that can be cultivated at low cost in simple media at a large scale, making it an attractive production platform for recombinant spike protein and other biopharmaceuticals in low-income countries.

## Introduction

In late 2019, the severe acute respiratory syndrome coronavirus 2 (SARS-CoV-2) was identified as the causative agent for the coronavirus disease 19 (COVID-19) ^1^. Since then, the virus is spreading throughout the world, causing not only humanitarian but also severe economic crisis. One of the most important proteins for the pathogenicity of the SARS-CoV-2 virus is the trimeric spike protein on its surface ^2–4^. This 1273-amino acids protein contains 22 N-linked glycans and only traces of O-linked glycans ^5^. Each spike protein monomer is built of two main subunits S1 and S2, with S1 mediating receptor binding and S2 membrane anchoring ^6–8^. Cleavage between the S1 and S2 subunits occurs during spike protein synthesis in the Golgi apparatus by a furin-like protease at a multi-basic cleavage site, leaving the S1 and S2 subunits linked by non-covalent interactions ^8, 9^. The S1 subunit contains the receptor binding domain (RBD), which binds to the cellular surface receptor angiotensin-converting enzyme 2 (ACE2) ^10–12^. Binding to ACE2 requires at least one RBD in the spike protein trimer to be in the “up” conformation. Accordingly, roughly half of the ~25 spike protein trimers present on the surface of a virion are in the “one RBD up” conformation ^2–4^. After binding of the RBD to the ACE2 receptor, structural changes occur in the spike protein trimer that expose the S2’ sites located immediately upstream of the fusion peptide, leading to cleavage by the transmembrane protease serine 2 (TMPRSS2) ^13, 14^. S2’ sites can also be cleaved by cathepsin L in the endosome after endocytosis of virions ^15^. Cleavage at S2’ leads to the shedding of S1 subunits and triggers a cascade of folding events resembling a “jackknife mechanism”, during which the fusion peptides are inserted into the target cell membrane. Further folding bends the viral and host membranes toward each other, leading to membrane fusion and viral entry ^8, 16^.

As the major protein on the virion surface, the spike protein is the prominent target of neutralizing antibodies and, therefore, all currently admitted vaccines use the spike protein as antigen ^16–19^. While mRNA- and vector-based vaccines have proven efficient and fast to establish, they do have side effects including heart inflammation and blood clots. Moreover, they are difficult to produce and to handle in lower-income countries ^17^. In contrast, recombinant protein based vaccines have less side effects and are easier to produce once their production platform is established ^20^. While potent neutralizing antibodies bind to the RBD, the RBD lacks other neutralizing epitopes present on the full-length spike protein ^16–18^. Hence, it appears advisable to produce the prefusion-stabilized, full-length spike protein for protein-based vaccines and for serological tests. Accordingly, a phase III clinical trial for the Novavax NVX-CoV2373 vaccine based on recombinant prefusion-stabilized full-length spike protein trimers has been completed recently with the vaccine showing 90% protection against COVID-19 ^21^. Because of its size and its 22 N-linked glycans, the production of full-length spike protein has only been achieved in mammalian and insect cells ^5–7, 10, 21–23^, while yeast, plant, and microalgal expression hosts have been limited to the production of spike protein fragments, mostly the RBD ^24–32^. Moreover, in the plant and microalgal expression hosts, the RBD was expressed as a cytosolic or ER-resident protein, requiring purification from whole cells ^24–27^. The development of simple production platforms for the secretion of complex therapeutic glycoproteins such as the full-length spike protein is desirable to facilitate their production in low-income countries. Microalgae appear to be a good choice here, as they can be grown at large scale in very simple media and do not produce toxic compounds ^33^. In general, glycosylation is essential for protein folding, stability, and functionality and is the most eminent post-translational modification in biopharmaceuticals ^34^.

Here we demonstrate the production and secretion of full-length SARS-CoV-2 spike protein in the unicellular green alga *Chlamydomonas reinhardtii*. We show that the protein can be enriched by ammonium sulfate precipitation from the culture medium and is functional, as judged by its ability to interact with human ACE2.

## Results

### Production and secretion of full-length SARS-CoV-2 spike protein in *Chlamdomonas reinhardtii*

To engineer Chlamydomonas toward the production and secretion of the full-length SARS-CoV-2 spike protein, we reverse translated the protein with optimal Chlamydomonas codon usage. We included the endogenous secretion peptide but excluded the C-terminal membrane anchor ^1^. Subsequently, we inserted introns of the Chlamydomonas RuBisCo small subunit gene *RBCS2* into the coding region, as proposed previously ^35, 36^. To facilitate gene synthesis, the sequence was divided into a 2610-bp and a 2851-bp fragment. Both fragments were flanked by BsaI recognition sites, following the syntax of the Chlamydomonas MoClo toolkit ^37^ for standardized level 0 parts in position B3 (*CoV-2 up*) and B4 (*CoV-2 down*) (Figure 1A). We first wanted to test whether Chlamydomonas can produce the SARS-CoV-2 spike protein and whether the endogenous secretion signal is sufficient to direct the protein for secretion into the culture medium. Following the single-step reaction assembly strategy for full transcriptional units, called level 1 modules ^37, 38^, we combined the strong, constitutive *HEAT SHOCK PROTEIN 70A-RBCS2 fusion* promoter, the two coding sequences *CoV-2 up* and *CoV-2 down*, an HA-tag coupled to an octa-histidine purification tag and the *RIBOSOMAL PROTEIN L23* terminator. Next, a level 2 multigene device was assembled using a level 1 module containing a spectinomycin resistance cassette (*aadA*) and the level 1 module for the expression of the SARS-CoV-2 spike protein (Figure 1B, WT-S). Finally, the level 2 device was transformed into the CC-4533 strain harboring an insertion in the *SIR2-LIKE NAD*(+) *DEPENDENT PROTEIN DEACETYLASE* (*SRTA*) gene. Chlamydomonas strains with defects in this gene facilitate the stable expression of foreign genes to high levels ^39, 40^. Immunoblot analyses of cell lysates from a spectinomycin resistant transformant revealed a protein migrating above the 130 kDa marker that was specifically detected by the HA antibody (Figure 2C). The calculated molecular mass for our SARS-CoV-2 spike protein design is 137 kDa. However, no signal was obtained for proteins from the culture medium. We therefore conclude that Chlamydomonas can produce the full-length recombinant spike protein but the endogenous secretion peptide cannot target the protein to the secretory pathway in Chlamydomonas.

**Figure 1.**
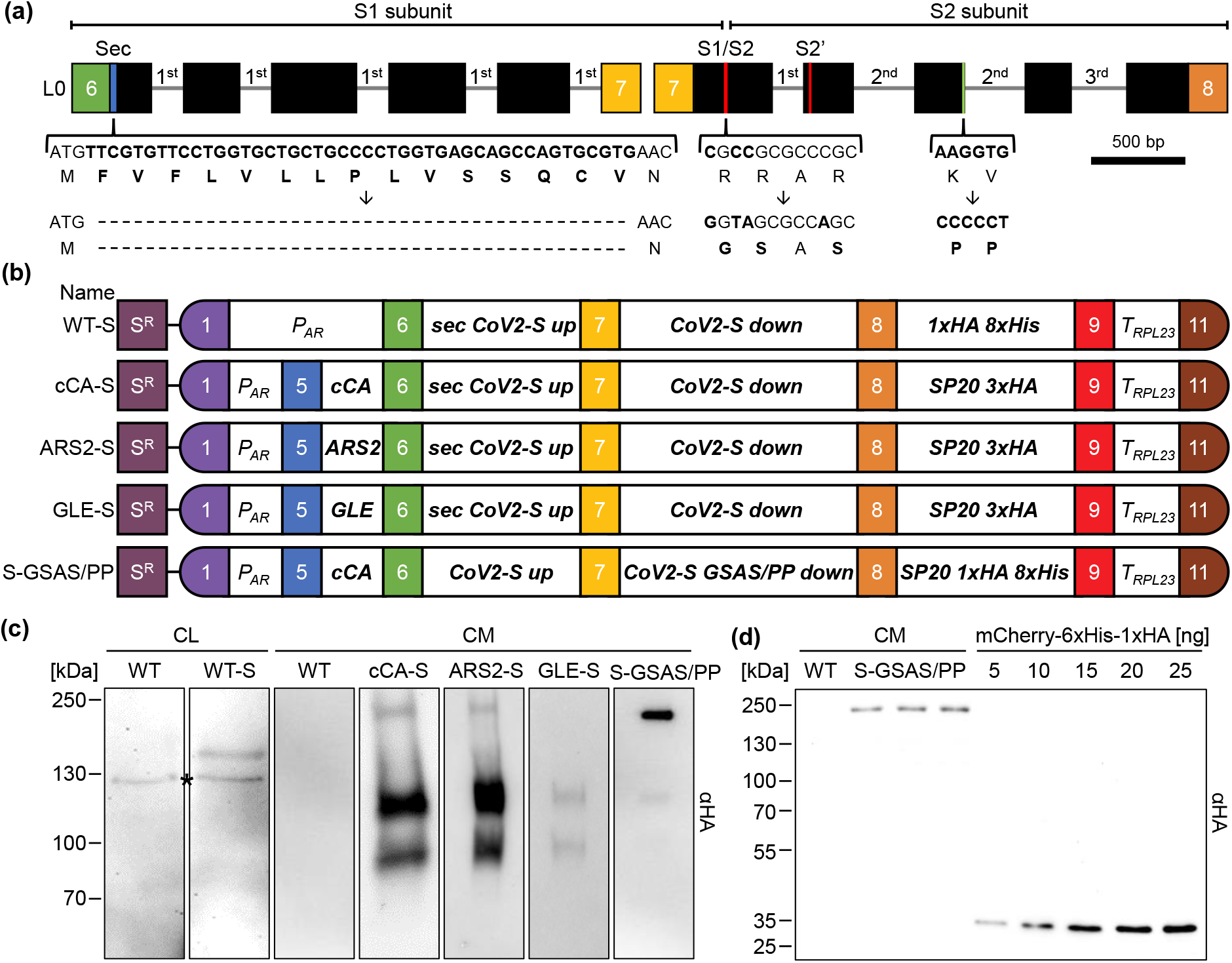
Design of constructs for the production and secretion of the SARS CoV-2 spike protein in Chlamydomonas. (a) Schematic representation of the two level 0 parts encoding the spike protein. The codon adapted coding region is depicted by black boxes and incorporated *RBSC2* introns by thin lines. The braces highlight regions modified by site directed mutagenesis and the resulting amino acid modifications. Colored boxes represent fusion sites according to the MoClo syntax. (b) Level 2 MoClo devices consisting of a level 1 module containing the *aadA* resistance marker under control of the *PSAD* promoter and terminator (S^R^), and level 1 transcriptional units for the expression of the spike protein. The latter contain the *HSP70A-RBCS2* promoter (*P_AR_*), sequences encoding various secretion signals (cCA, ARS, GLE, sec), the two parts of the spike protein (CoV2-S up and down), a glycomodule of 20 serine/proline repeats (SP20) fused to the HA epitope and/or an octa-his tag, and the *RPL23* terminator (*T_RPL23_*). (c) Production and secretion of the spike protein in transformants generated with the five level 2 devices listed in (b). Shown are representative immunoblots detecting HA-tagged proteins in cell lysates (CL) (corresponding to 2 μg chlorophyll) and 1.7 mL of culture medium (CM) after TCA precipitation. The asterisk indicates a cross reaction of the HA antibody. (d) Three independent preparations of spike protein secreted by a S-GSAS/PP transformant into 1.7 mL culture medium (CM) and precipitated with TCA, and increasing amounts of purified recombinant mCherry carrying a C-terminal HA tag were analyzed by immunoblotting using an HA antibody.

**Figure 2.**
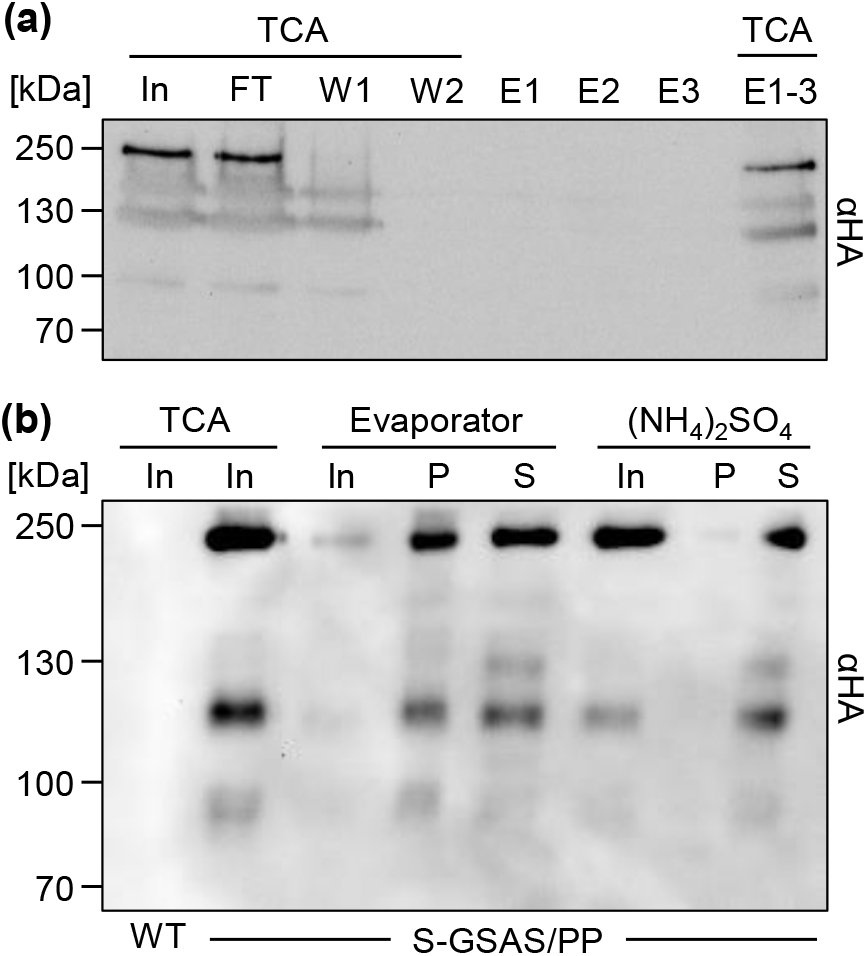
Enrichment of secreted SARS-CoV2 spike protein from Chlamydomonas culture medium. (a) Attempted purification of the octa-histidine-tagged spike protein from culture medium. 250 mL culture medium from a S-GSAS/PP transformant was applied to a nickel-NTA affinity column. The column was washed twice and bound proteins eluted with imidazole in three fractions. Proteins in 1.7 mL of input (In), flow-through (FT), and washes (W1 and W2), and those in the entire eluate (E1-3) were precipitated with TCA. Precipitates and 20 μL of three elution fractions (E) were separated on an 8% SDS-polyacrylamide gel and analyzed by immunoblotting using an HA antibody. (b) Comparison of concentration methods for the spike protein. Culture medium of Chlamydomonas wild-type cells (WT) and of a S-GSAS/PP transformant was precipitated with TCA (In - theoretical concentration factor (CF): 85x). Proteins in 100 mL S-GSAS/PP culture medium were concentrated 17x with a rotary evaporator (Evaporator - In) followed by two concentration steps with centrifugal filters (MWCO 3000 Da, theoretical CF: 34x). The concentrate was then centrifuged to pellet aggregated proteins (Evaporator - P) and have soluble proteins in the supernatant (Evaporator - S). Proteins in 100 mL S-GSAS/PP culture medium were precipitated with ammonium sulfate (NH_4_SO_4_ - In, theoretical CF: 34x). Resuspended proteins were centrifuged to pellet aggregated proteins (NH_4_SO_4_ - P) and have soluble proteins in the supernatant (NH_4_SO_4_ - S). All proteins were separated on an 8% SDS-polyacrylamide gel and analyzed by immunoblotting using an HA antibody

To enable secretion of the spike protein, the MoClo level 1 module was reconstructed by adding the coding regions for three different N-terminal secretion signals of native extracellular Chlamydomonas proteins, namely carbonic anhydrase (cCA) ^41^, arylsulfatase 2 (ARS2) ^42^ and gamete lytic enzyme (GLE) ^43^. In the following, the constructs are named by the secretion signal (cCA-S, ARS2-S, and GLE-S). Additionally, the coding region for a C-terminal synthetic glycomodule of 20 serine/proline repeats was added, which was reported to enhance secretion yields by up to 12-fold in Chlamydomonas ^43^. To increase the detectability of the spike protein, the glycomodule was equipped with a triple HA motif (SP20 3xHA). As before, the resulting level 1 modules were assembled into level 2 devices with the *aadA* cassette. Proteins in the culture medium of various spectinomycin resistant transformants were precipitated and analyzed by immunoblotting. We noticed that the chemiluminescence signal of secreted spike protein was much stronger in transformants generated with the cCA-S and ARS2-S constructs when compared with transformants generated with GLE-S (Figure 2C), pointing to a more efficient secretion via the cCA and ARS secretion peptides. Moreover, the HA antibody specifically detected three protein bands in the culture medium of cCA-S and ARS2-S transformants with apparent molecular masses of ~240, ~120 and ~90 kDa. The calculated molecular mass of the mature spike protein lacking the secretion signal is 142 kDa, which does not fit to any of the detected bands. It has been shown that the synthetic SP20 glycomodule is efficiently glycosylated during its passage through the secretory pathway in Chlamydomonas ^43^. Moreover, the spike protein contains 22 N-linked glycans ^5^. Hence, it appears likely that glycosylation of the SP20 module and the spike protein itself takes place and that the protein migrating at ~240 kDa represents the glycosylated full-length protein. The latter appears to be efficiently processed, giving rise to two C-terminal fragments, accounting for the weak signal for the full-length ~240 kDa protein and the strong signals at ~120 and ~90 kDa. In the weakly expressing GLE-S transformants, the full-length protein appears to be below the detection limit and only the two C-terminal cleavage products are detected (Figure 2C).

During its synthesis, the SARS-CoV-2 spike protein is cleaved into two functional subunits S1 and S2 by a furin-like protease in the Golgi apparatus ^8, 9^ (Figure 1A). Cleavage at the S1/S2 boundary is strongly enhanced in the SARS-CoV-2 spike protein by the presence of a “RRAR” furin cleavage site that is missing in the SARS-CoV spike protein ^6, 7, 9, 10, 44^. We wondered, whether processing of the SARS-CoV-2 spike protein produced in Chlamydomonas was also mediated by a furin-like protease. To test this, we reconstructed the level 0 parts of the coding sequence. At first, we removed sequences coding for the N-terminal endogenous signal peptide, since it was unable to mediate secretion in Chlamydomonas (Figure 1C). Next, we exchanged the sequences encoding the furin cleavage site (S_682_RRAR↓S_685_) via site directed mutagenesis by nucleotides coding for S_682_GSAS S_685_, which is supposed to completely inactivate the furin cleavage site ^44^. Furthermore, we substituted codons for K_986_V_987_ situated at the beginning of the central helix by two codons for proline, which was reported to increase expression yields and to stabilize the prefusion conformation of coronavirus spike proteins resulting in higher immunogenicity ^7, 45, 46^. Finally, we equipped this new variant of the CoV-2 spike protein with a SP20 glycomodule carrying a single HA tag and an octa-histidine tag. We employed only the cCA secretion signal, as it proved to be efficient (construct S-GSAS/PP in Figure 1A and B). Immunoblot analyses of proteins precipitated from the culture medium of a S-GSAS/PP transformant with the HA antibody revealed a prominent protein band at ~240 kDa and very minor ones at ~120 kDa and ~90 kDa (Figure 1C). Hence, eliminating the furin cleavage site appears to almost completely abolish proteolytic cleavage of the spike protein during its passage through the secretory pathway, as has been observed in human cells ^6, 10, 44^.

To estimate the amounts of full-length spike protein secreted into the culture medium by the S-GSAS/PP transformant, we performed quantitative immunoblotting. To this end, proteins in the medium of S-GSAS/PP cultures grown for five days after inoculation were precipitated and separated on an SDS-gel next to known amounts of recombinant mCherry carrying a C-terminal HA tag (Figure 1D). Immunodetection with the HA antibody and quantification of the signals revealed an average concentration of full-length spike protein of 11.2 ± 1.8 μg/L (n = 3, ± SD).

### SARS-CoV-2 spike protein from Chlamydomonas cannot be purified by IMAC but is readily concentrated by ammonium sulfate precipitation

After the successful production of full-length SARS-CoV-2 spike protein in Chlamydomonas, we aimed to purify the secreted protein from culture medium of a S-GSAS/PP transformant via immobilized metal affinity chromatography, taking advantage of the C-terminal octa-histidine tag. However, the spike protein failed to bind to the nickel resin and largely remained in the flow through (Figure 2A). The same result was obtained when the medium was supplied with 2% Triton or 250 mM NaCl, with longer incubation times, or when we used cobalt beads instead of nickel beads. An affinity purification with immobilized HA antibodies also failed (not shown). To enrich the protein by other means, we concentrated proteins in the culture medium with a rotary evaporator, followed by two runs through centrifugal filters. Although an enrichment of the spike protein was achieved, about half of it was found in aggregates (Figure 2B). As an alternative, we precipitated proteins in the culture medium with ammonium sulfate and could recover virtually all of the spike protein, with very little found in aggregates (Figure 2B). Hence, ammonium sulfate precipitation appears most suitable for enrichment.

### SARS-CoV-2 spike protein from Chlamydomonas binds to the hACE2 receptor

Next, we wanted to test the functionality of the spike protein produced in Chlamydomonas. The ability of the protein to bind to the human ACE2 receptor was described previously ^10–12^ and binding to ACE2 is an accepted assay to verify the functionality of recombinant spike protein ^29^. We incubated proteins enriched by ammonium sulfate precipitation from a S-GSAS/PP culture with or without recombinant hACE2 containing a C-terminal deca-histidine tag. Nickel resin was added and bound proteins eluted. As shown in Figure 3A, full-length spike protein co-eluted with hACE2. No spike protein was eluted if hACE2 was absent. We noticed that less spike protein was recovered with hACE2 than supplied in the input, indicating that only part of the supplied spike protein bound to the hACE2 receptor. Moreover, unbound spike protein in the supernatant was almost completely converted to the ~90-kDa breakdown product, presumably during the 30-min incubation at 37°C. As a control, we produced the SARS-CoV-2 RBD fused to sfGFP in transiently transfected human 293T cells. Secreted proteins were concentrated with centrifugal filters and incubated with recombinant hACE2. As expected, we observed the RBD-sfGFP fusion protein to co-elute with hACE2 (Figure 3B). Hence, the *in vitro* pull-down assay works equally well with human-derived RBD and Chlamydomonas-derived full-length spike protein.

**Figure 3.**
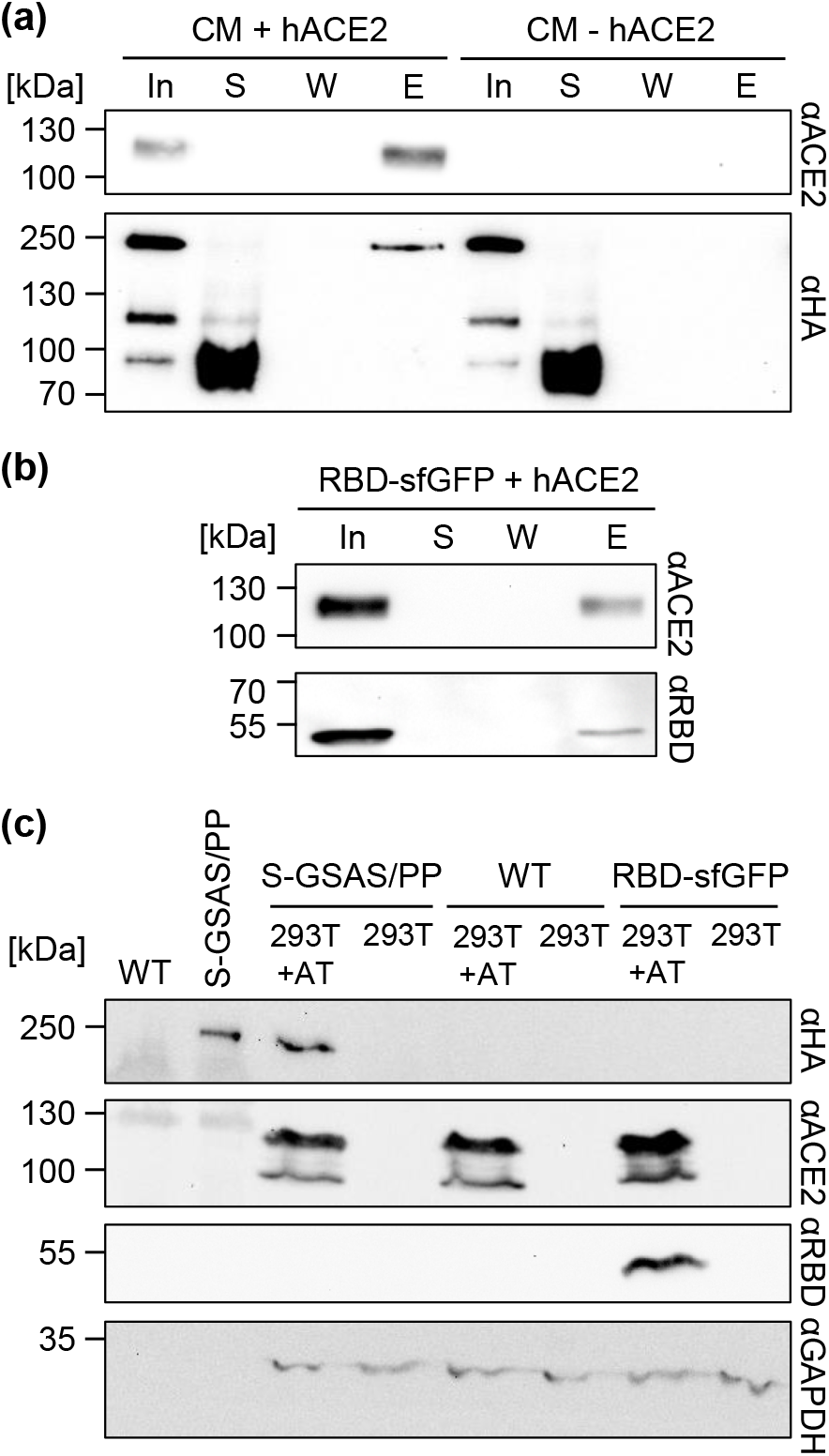
Assays for testing the functionality of SARS-CoV2 spike protein produced in Chlamydomonas. (a) *In vitro* pull-down assay with full-length spike protein from a S-GSAS/PP transformant and recombinant hACE2. Proteins from the culture medium (CM) of a S-GSAS/PP transformant that had been precipitated with ammonium sulfate and dialyzed against PBS were incubated in the presence or absence of recombinant hACE2 containing a deca-histidine tag. Nickel resin was added, sedimented by centrifugation and the supernatant (S) collected. The resin was washed once (W), and bound proteins eluted with imidazole (E). Proteins in S, W and E were precipitated with TCA (100%) and analyzed next to 10% of the input (In) by immunoblotting using antibodies against hACE2 and the HA epitope. Shown is one representative experiment out of three with similar outcome. (b) *In vitro* pull-down assay with RBD-sfGFP and recombinant hACE2. The experiment was conducted as in (a) but using human derived RBD-sfGFP concentrated with centrifugal filters. (c) Cell-based functionality assay with full-length spike protein from a S-GSAS/PP transformant and RBD-sfGFP. Proteins in culture medium from a S-GSAS/PP transformant and from wild type were concentrated by ammonium sulfate precipitation, dialyzed against PBS, and applied to 293T cells constitutively overexpressing h<u>A</u>CE2 and h<u>T</u>MPRSS2 (293T+AT) and regular 293T cells. As control, RBD-sfGFP concentrated with centrifugal filters was applied. Cells were washed, lysed and total proteins subjected to SDS-PAGE and immunoblot analysis using antibodies against the HA epitope, ACE2, and the RBD. GAPDH was detected as endogenous housekeeping control. Shown is one representative experiment out of four with similar outcome.

To complement the *in vitro* binding assay, we performed a second functionality assay based on the binding of spike protein to hACE2 on human cells. For this, we concentrated secreted proteins from culture medium of a S-GSAS/PP transformant by ammonium sulfate precipitation. Concentrated proteins were then applied to 293T cells constitutively overexpressing h<u>A</u>CE2 and h<u>T</u>MPRSS2 (293T+AT) and as a control to regular 293T cells. Importantly, we did not observe any cytotoxic or other negative effects of this treatment on the cells by microscopy. After washing, cells were lysed and proteins in the lysates analysed by SDS-PAGE and immunoblotting. As shown in Figure 3C, the spike protein was detected in lysates from cells expressing hACE2, while it was not detected in lysates from cells lacking hACE2. Accordingly, when the experiment was conducted with concentrated recombinant RBD-sfGFP, a signal for RBD was obtained only in lysates from cells expressing hACE2 (Figure 3C). No signal at the migration position of the spike protein in SDS-gels was obtained if cells were treated with concentrated secreted proteins from wild-type cells (Figure 3C) Taken together, these results provide strong evidence that full-length SARS-CoV-2 spike protein produced in Chlamydomonas is functional.

## Discussion

Here we report on the construction of Chlamydomonas strains that secrete functional full-length SARS-CoV-2 spike protein into the medium. To our knowledge, this has only been achieved with mammalian and insect cell-based expression systems. Recent reports on the expression of SARS-CoV2 spike protein in Chlamydomonas were limited to intracellularly expressed RBD ^24, 26^. The advantage of Chlamydomonas is that it is a GRAS organism that can be cultivated at low cost in simple media at a large scale, making Chlamydomonas an attractive production platform for human therapeutical proteins in low-income countries ^33, 47^.

During our endeavor to produce the spike protein in Chlamydomonas, we have learned that its native signal peptide does not guide the protein to the secretory pathway in Chlamydomonas. For this, a signal peptide of a native Chlamydomonas protein was required, with those from cCA and ARS2 being more efficient than that of GLE (Figure 1C). We observed that during its passage through the secretory pathway the spike protein is processed at the multi-basic furin cleavage site, just like in mammalian cells ^6, 10, 44^, which argues for the presence of a furin-like protease in the ER/Golgi system in Chlamydomonas. An O-glycosylation was reported at T678 close to the furin cleavage site with a potential role in regulating cleavage ^48^. If so, the efficient cleavage observed in Chlamydomonas might indicate that this O-glycosylation is realized in Chlamydomonas.

A C-terminal glycomodule of 20 serine-proline repeats (SP20) apparently was required for efficient spike protein secretion, as we failed to identify transformants secreting the protein if it did not contain the SP20 module (not shown), corroborating earlier findings on the stimulating effect of the SP20 module on protein secretion in Chlamydomonas ^43^. Purification of the spike protein via an HA or an octa-histidine tag at the very C-terminus failed, albeit we could detect the HA tag in immunoblots (Figures 1A and 2A). Possible reasons for the unsuccessful affinity purification could be (i) proteolytic removal of the tag during its passage through the secretory pathway or in the culture medium; (ii) steric hindrance by unfavorable folding of the tag; (iii) steric hindrance by glycans at the SP20 module immediately upstream of the tag; (iv) interference by other secreted proteins.

Concentration of secreted proteins by a rotatory evaporator and centrifugal filters led to aggregation of part of the spike protein, while enrichment by ammonium sulfate precipitation resulted in soluble spike protein (Figure 2B). Functionality of the Chlamydomonas-produced spike protein was deduced from its ability to interact with soluble recombinant ACE2 and ACE2 expressed on human cells (Figure 3). Since glycosylation has been shown to be essential for the correct folding of soluble and functional spike protein ^29, 49^, it appears that N-glycosylation realized in Chlamydomonas promotes proper spike protein folding. Also, the much larger apparent than calculated molecular mass of the secreted versus the cytosolically produced spike protein point to successful glycosylation beyond that on the SP20 module (Figure 1). Hence, our results are in line with recent reports on the production of functional forms of human vascular endothelial growth factor ^50^, human epithelia growth factor ^51^, and RBD ^24^ in Chlamydomonas, indicating that Chlamydomonas is a suitable production platform for human therapeutic proteins. Nevertheless, it is important to keep in mind that N-glycosylation patterns realized by Chlamydomonas on secreted proteins are distinct from those realized in human cells, with glycoengineering of Chlamydomonas toward a more human-like glycosylation pattern representing an option for the future ^52^.

The assembly and testing of the constructs for expressing the various spike proteins could be realized rapidly in iterative cycles thanks to the MoClo strategy, easy transformation and short generation times of Chlamydomonas, and the use of expression strains ^36–39^. Clearly, these features will facilitate adaptations to SARS-CoV-2 mutant variants and help addressing the remaining bottlenecks with spike protein production in Chlamydomonas: first, we need to find a method for purifying the protein from the culture medium, *e.g*., by changing the position of tags to the N-terminus, or by employing other tags like GST, MBP, FLAG, or StrepII. Second, we need to enhance yields. By quantitative immunoblotting we estimated a yield for secreted spike protein of 11.2 μg/L (Figure 1D). Because the large, glycosylated spike protein will be less efficiently transferred to nitrocellulose by semi-dry blotting than the small non-glycosylated mCherry used as standard, we are definitely underestimating yields. LC-MS/MS-based methods based *e.g*. on QconCATs ^53^ should be employed in future. Reported yields of secreted proteins produced in Chlamydomonas were 100 μg/L for human erythropoietin ^42^, 28 μg/L for human vascular endothelial growth factor ^50^, 100 μg/L for a human epithelial growth factor fusion with luciferase ^51^, 0.7 mg/L for various reporter proteins ^54^, 15 mg/L for mVenus ^43^, and 10-12 mg/L for luciferase alone or fused to an ice-binding protein ^41, 55^. Hence, our yields are in the same range as those reported for other human therapeutic proteins, but far below those reported for the simpler reporter proteins. A potential factor limiting yield might be the efficiency of N-glycosylation and associated protein folding, which must be a challenge for the spike protein given its large size and 22 N-linked glycans ^5^. Accordingly, the ER chaperone BiP was found as a major contaminant of spike protein preparations ^8^. Therefore, enhanced expression of human proteins in Chlamydomonas might be achieved by the overexpression of components of the ER protein folding machinery, or of the bZIP1 transcription factor regulating the Chlamydomonas ER unfolded protein response ^56, 57^. Moreover, protein secretion yields in Chlamydomonas can be strongly improved by optimizing culture conditions ^55, 58^.

## Methods

### Strains and Culture conditions

*Chlamydomonas reinhardtii* strain CC-4533 harboring an insertion in the *SRTA* gene (LMJ.RY0402.148523) ^39^ was obtained from the Chlamydomonas Resource Center (https://www.chlamycollection.org/). Cells were grown in 10-mL cultures in Tris-Acetate-Phosphate (TAP) medium ^59^ on a rotatory shaker at a constant light intensity of ~40 μmol photons m^−2^ s^−1^. Transformation was performed with the glass beads method as described previously ^60^ with constructs linearized with NotI. Transformants were selected on 100 μg/mL spectinomycin.

To validate the functionality of SARS-CoV2 S-protein, HE293T cells that stably overexpress h<u>A</u>CE2 and h<u>T</u>MPRSS2 (293T+AT; CL0015 VectorBuilder) and non-transfected HEK293T (293T; ATCC, CRL-3216™) cells were used. The cells were maintained in Dulbecco’s Modified Eagle’s Medium (DMEM) supplemented with 10 % fetal calf serum and antibiotics (500 μg/mL neomycin, 7.5 μg/mL blasticidin, 1.5 μg/mL puromycin for 293T+AT and 65 μg/mL penicillin G, 100 μg/mL streptomycin sulfate for 293T) in humified atmosphere at 37 °C and 5% CO_2_.

### Expression of RBD-sfGFP

As a control for the activity assays, RBD-sfGFP was expressed in 293T cells. The cells were seeded into 175 cm^2^ flasks and incubated until they reached 60 % confluence. Transfection was performed using polyethyleneimine (branched, 408727, Sigma-Aldrich). For this, 45 μg of plasmid DNA (pcDNA3-SARS-CoV-2-S-RBD-sfGFP ^61^, http://n2t.net/addgene:141184, kindly provided by Erik Procko) were incubated with 2 mL of DMEM. After 5 min, 805 μL polyethyleneimine (1 mg/mL) was added, immediately mixed and incubated for 10 min. The culture medium was removed from the cells and the mixture was then carefully added. After two hours, the medium was replaced by fresh serum-free medium without phenol red (P04-710629, PAN Biotech). Expression took place for three days. The culture medium containing RBD-sfGFP was then collected and centrifuged (1,000 g, 10 min, 4°C), and concentrated with centrifugal filters (MWCO 10 kDa; Amicon^®^ Ultra-15).

### Cloning

The 1212-amino acids N-terminal portion of the SARS-CoV-2 spike protein) lacking the membrane anchor (UniProt: P0DTC2) was reverse translated using the most commonly used Chlamydomonas codons. For stable foreign gene expression ^36^, the first *Chlamydomonas RBCS2* intron was inserted six times with the flanking sites GAG/intron/G, the second intron was inserted twice with flanking sites GCG/intron/GC, and the third was inserted once using ACG/intron/G as flanking sites. A single internal BsaI recognition site was removed by changing the used codon for threonine (ACC) into ACG. The sequence was split into two fragments, which were flanked with BsaI recognition sites such that, upon BsaI digestion, fragments with AATG and AGGT or AGGT and TTCG overhangs are generated for the B3 and B4 positions of level 0 parts according to the MoClo syntax for plant genes ^38^. Synthesis and cloning into the pUC57 vector was done by BioCat (Heidelberg), resulting in level 0 parts pMBS704 (*secCoV2-up*) and pMBS705 (*CoV2-down*). To remove the sequence coding for the N-terminal endogenous secretion signal, pMBS704 was used as template for PCR using primers 5’-AAAGAAGACAAAATGAACCTGACCACCCGCACCCAGC-3’ and 5’-AAAGAAGACTTCATTTGAGACCTTTATATCTAGATG-3’. The resulting 5170-bp PCR product was digested with BbsI and assembled by ligation, giving rise to level 0 part pMBS706 (*CoV2-up*). pMBS705 was used as template for site directed mutagenesis PCR to replace codons for the furin recognition site (S1/S2) “RRAR” with codons for “GSAS”, as described previously ^7, 44^ and to replace the codons for “KV” with two proline codons, as described previously ^45, 46^. The following primer combinations were chosen to amplify three fragments (overhangs are underlined, lower case letters indicate base changes): 5’-AAAGAAGACGT<u>CTCA</u>AGGTGCCCGTGGCCA-3’ and 5’-AAAGAAGACTT<u>taCc</u>GGGGCTGTTGGTCTGGGTCTGG-3’ (218 bp), 5’-AAAGAAGACAA<u>gGta</u>GCGCtaGCAGCGTGGCCAGCCAGAGCA-3’ and 5’-AAAGAAGACTT<u>GTCG</u>TTCAGCACGCTGCTGATGGC-3’ (1390 bp), and 5’-AAAGAAGACAA<u>CGAC</u>ATCCTGAGCCGCCTGGACcccccGGAGGTGAGCTTGC-3’ and 5’-AAAGAAGACTT<u>CTCG</u>CGAACCCCACTTGATGTACT-3’ (1302 bp). The three PCR products were combined with pAGM9121 ^38^, digested with BbsI and directionally assembled by ligation into the level 0 part pMBS708 (*CoV2-GSAS/PP-down*). A level 0 part encoding the arylsulfatase 2 secretion peptide (*ARS2*) was generated via the annealing of oligonucleotides 5’-GAAGACAA<u>CCAT</u>GAGCCTGGCCACCCGCCGCTTCGGCGCCGCCGCCGCCCTG CTGGTGGCCGCCTGCGTGCTGTGCACCGCCCCCGCCTGGGC<u>AATG</u>AAGTCTTC-3’ and 5’-GAAGACTTCATTGCCCAGGCGGGGGCGGTGCACAGCACGCAGGCGGCCACCAGCAGGGCGGCGGCGGCGCCGAAGCGGCGGGTGGCCAGGCTCATGGTTGTCTTC-3’. The product was combined with destination vector pAGM1276 ^38^, digested with BbsI and ligated to yield pMBS489 (*spARS2*). The sequence encoding the secretion signal of the gamete lytic enzyme (*GLE*) was similarly produced via the annealing of oligonucleotides 5’-GAAGACAACCATGAGCCTGGCCACCCGCCGCTTCGGCGCCGCCGCCGCCCTG CTGGTGGCCGCCTGCGTGCTGTGCACCGCCCCCGCCTGGGC<u>AATG</u>AAGTCTTC-3’ and 5’-GAAGACTT<u>CATT</u>GCCCAGGCGGGGGCGGTGCACAGCACGCAGGCGGCCACCA GCAGGGCGGCGGCGGCGCCGAAGCGGCGGGTGGCCAGGCTC<u>ATGG</u>TTGTCTT C-3’ followed by digestion with BbsI and ligation with destination vector pAGM1276 ^38^, giving rise to level 0 part pMBS490 (*spGLE*). A level 0 part encoding the hemagglutinin (HA) motif, RGS linker and an oct-histidine tag was obtained via the annealing of oligonucleotides 5’-AAGAAGACAA<u>TTCG</u>TCTGGTTACCCCTACGACGTGCCCGACTACGCTCGCGGC AGCCACCACCACCACCACCACCACCACTAA<u>GCTT</u>AAGTCTTCAA-3’ and 5’-TTGAAGACTT<u>AAGC</u>TTAGTGGTGGTGGTGGTGGTGGTGGTGGCTGCCGCGAGC GTAGTCGGGCACGTCGTAGGGGTAACCAGA<u>CGAA</u>TTGTCTTCTT-3’. Digestion of the annealing product and the destination vector pAGM1301 with BbsI and ligation yielded pMBS723 (*1xHA-8xHis*). The SP20 glycomodule ^43^ coupled to a triple HA motif was reverse translated using the most commonly used Chlamydomonas codons and flanked with BsaI recognition sites, producing overhangs for the B5 position according to the MoClo syntax for plants ^38^. Synthesis and cloning into the pUC57 vector were done by BioCat (Heidelberg), yielding level 0 part pMBS514 (*SP20-3xHA*). The plasmid pMBS514 was used as template to replace the sequence encoding two HA motifs by a sequence encoding the RGS-8xHis motif by PCR using primers 5’-TAAGCTTTGAGACCTTCCAATATC-3’ and 5’-TCATTAGTGGTGGTGGTGGTGGTGGTGGTGGCTACCGCGAGCGTAGTCGGGCACGTCGTA-3’. The 2810-bp PCR product was phosphorylated with polynucleotide kinase (NEB) and circularized with T4 DNA ligase (NEB) as descripted previously ^62^, generating pMBS659 (*SP20-1xHA-8His*). All PCRs were conducted with Q5 High-Fidelity Polymerase (NEB) following the manufacturer’s instructions. Sequences of the level 0 parts were verified by Sanger sequencing (Seqlab). Level 1 module assembly was performed by combining newly constructed level 0 parts with level 0 parts (pCM) from the Chlamydomonas MoClo toolkit ^37^ and the respective destination vector pICH47742 ^38^. For directionally assembly, the following level 0 parts were chosen: A1-B1-pCM0-015 (*HSP70A-RBCS2* promoter + 5’-untranslated region (UTR)); A1-B2-pCM0-017 (*HSP70A-RBCS2* promoter + 5’-UTR); B2-pCM0-051 (*sp carbonic anhydrase (cCA*)); B2-pMBS489 (spARS); B2-pMBS490 (*spGLE*); B3-pMBS704 (*CoV2-up*); B2-B3-pMBS706 (*sec-CoV2-up*); B4-pMBS705 (*CoV2-down*); B4-pMBS708 (*CoV2-GSAS/PP-down*); B5-pMBS723 (*1xHA-8xHis*); B5-pMBS514 (*SP20-3xHA*); B5-pMBS659 (*SP20-1xHA-8His*); B6-C1-pCM0-119 (*RPL23 3*’-UTR). The individual level 0 parts were released by BsaI digestion and assembled into five level 1 modules using T4 DNA ligase according to Figure 1B. The level 1 modules were then combined with pCM1-01 (level 1 module with the *aadA* gene conferring resistance to spectinomycin flanked by the *PSAD* promoter and terminator) from the Chlamydomonas MoClo kit ^37^, with plasmid pICH41744 containing the proper end-linker, and with destination vector pAGM4673 ^38^, digested with BpiI, and ligated to yield the five level 2 devices displayed in Figure 1B.

### Cloning and purification of recombinant mCherry-6His-1xHA

The mCherry coding sequence was amplified via PCR using plasmid CD3-960 ^63^ as template and primers 5’-AAAAGCTAGCAAAGAATTCATGGTGAGCAAGGGCGAGGAG-3’ and 5’-TTTCTCGAGTTTGTACAGCTCGT<u>CCAT</u>GC-3’ flanked by NheI and XhoI recognition sites. The PCR product and pET21b were digested with NheI and XhoI, gel-purified, and ligated to get pET-mCherry-6His (pMS1034). This plasmid was used as a template for PCR to add the coding sequence for the 1xHA motif via the primers 5’-AAAA<u>GGATCC</u>TATCCGTATGATGTGCCAGATTATGCCTGAGATCCGGCTGCTAA CAAA-3’ and 5’-TTTT<u>GGATCC</u>GTGGTGGTGGTGGTGGTGCTCGAG-3’ (BamHI sites are underlined). The resulting 6140-bp PCR product was digested with BamHI and recircularized, giving rise to pET21b-mCherry-6xHis-1xHA (pMS1090). mCherry-6xHis-1xHA was expressed in ER2566 cells in LB-medium (100 μg/mL ampicillin) after inducing expression with 0.5 mM IPTG at 37 °C for 4 hours and purified under denaturing conditions as described previously ^53^ using nickel-charged resin. The concentration of recombinant mCherry-6His-1xHA was determined using the Pierce BCA protein assay kit following the manufacturer’s instructions.

### Protein analyses by SDS-PAGE

For whole-cell protein extraction, cells were pelleted and resuspended in 50 mM DTT, 50 mM Na_2_CO_3_, 2.5 % (w/v) SDS and 15 % (w/v) sucrose, boiled for 2 min at 95 °C and centrifuged. After determination of chlorophyll content ^64^, a sample volume corresponding to 2 μg chlorophyll was subjected to SDS-PAGE. For analysing proteins secreted into the culture medium, cells were grown to stationary phase. Cells in 1.7 mL culture were pelleted and the supernatant was transferred to a 2-mL reaction tube. Trichloroacetic acid (TCA) was added to a final concentration of 10%. The samples were incubated for 30 min on ice and centrifuged at 15,000 g for 15 min at 4 °C. The pellet was washed with 500 μL ice-cold acetone, resuspended in 20 μL of 75 mM Tris-HCL pH 6.8, 2.5 % (w/v) SDS, 100 mM DTT and 10 % (v/v) glycerol, boiled for 2-5 min at 95 °C and subjected to SDS-PAGE. After semidry blotting, immunodetection was performed by enhanced chemiluminescence using an INTAS imaging system. Primary antibodies used for immunodetection were mouse anti-HA (H9658, Sigma-Aldrich, 1:10,000), mouse anti-RBD (MAB105401, R&D Systems, 1:2,000) and mouse anti-ACE2 (sc-390851, Santa Cruz Biotechnology, 1:2,000). The secondary antibody was m-IgGκ BP-HRP (sc-516102, Santa Cruz Biotechnology, 1:10,000). Densitometric band quantifications after immunodetections were done with the FUS and IONCapt Advance program (PEQLAB).

### Nickel-nitrilotriacetic acid purification of SARS CoV-2 spike protein from culture medium

250 mL of culture medium of a S-GSAS/PP transformant grown to stationary phase were run over an affinity column containing 500 μL nickel-charged resin equilibrated with PBS (137 mM NaCl, 2.7 mM KCl, 81 mM Na_2_HPO_4_, 18 mM KH_2_PO_4_, pH 7.6). After two washing steps with 5 mL PBS containing 5 mM imidazole, bound proteins were eluted with 1.5 mL PBS containing 500 mM imidazole and three fractions of 0.5 mL each were collected.

### Ammonium sulfate precipitation of secreted proteins

Cells from 200 mL of culture of a S-GSAS/PP transformant grown to stationary phase were removed by three subsequent centrifugations at 4000 g for 5 min at 25 °C. The supernatant was transferred to a 500-mL beaker and grinded ammonium sulfate was slowly added under continuous stirring until the solution was saturated (106.6 g/200 ml). Stirring was continued for another 30 min. Precipitated proteins were pelleted by centrifugation at 4000 g for 45 min at 4 °C and the supernatant was carefully removed. The pellet was resuspended in 5 mL PBS and dialyzed (ZelluTrans 3.5 MWCO, Carl Roth) overnight against PBS. Finally, 1x protease inhibitor (cOmplete™, EDTA-free Protease Inhibitor Cocktail, Roche) was added to the solution according to the manufacturer’s instructions.

### SARS-CoV-2 spike protein binding assay to recombinant hACE2

Recombinant hACE2 with a C-terminal deca-histidine tag (SAE0064, Sigma-Aldrich) was resuspended in PBS to a final concentration of 50 ng/μL. 1 μg of hACE2 was mixed with 300 μL of proteins recovered after ammonium sulfate precipitation from a S-GSAS/PP culture and incubated for 30 min at 37 °C. The protein mixture was filled up to 1 mL with PBS, and 66 μL of nickel affinity resin (equilibrated with PBS) and 1x cOmplete™, EDTA-free Protease Inhibitor Cocktail (Roche) were added. After an incubation for 30 min on a circular rotator, samples were cooled for 10 min on ice, centrifuged at 100 g for 30 s at 4 °C and washed with 1 mL PBS containing 5 mM imidazole. Elution of bound proteins was performed with 50 μL PBS containing 500 mM imidazole. The resin was pelleted and the supernatant recovered. Samples from all steps were collected for further analysis. Flow-through, washing and elution fractions were TCA precipitated as described above.

### SARS-CoV 2 spike protein binding assay to cell-based hACE2

293T+AT and 293T cells were seeded into 6-well plates and grown until they reached 70% confluence. The medium was partially removed and 500 μL of proteins, recovered after ammonium sulfate precipitation from a S-GSAS/PP culture, were applied. After 1 h incubation at 37 °C and 5% CO_2_, the medium was removed and cells gently resuspended in PBS. After centrifugation at 800 g for 10 min at 4 °C, the supernatant was removed and the cell pellet lysed in ice-cold RIPA buffer (150 mM NaCl, 50 mM Tris pH 7.4, 1% Nonidet P-40, 0.1% SDS, 0.5% Sodium deoxycholate, 5 mM EDTA) containing 1x protease inhibitor cocktail (cOmplete™, EDTA-free Protease Inhibitor Cocktail, Roche). Cell debris was removed by centrifugation (8,000 g, 10 min, 4 °C). For immunoblot analysis, the lysates were separated on an 8% SDS-polyacrylamide gel and transferred to a nitrocellulose membrane. The membranes were blocked and incubated with specific antibodies: HA-tagged spike protein was detected with anti-HA antibody (H9658, Sigma-Aldrich, 1:10,000), hACE2 was detected with anti-ACE2 antibody (sc-390851, Santa Cruz Biotechnology, 1:2,000), RBD was detected using anti-RBD antibody (MAB105401, R&D Systems, 1:2,000). m-IgGκ BP-HRP was used as secondary antibody for detection (sc-516102, Santa Cruz Biotechnology, 1:10,000).

## Acknowledgements

We thank Benedikt Venn for helpful comments to the manuscript.

## Author contributions

A.K., J.N. and A.P. generated all constructs and performed all experiments. A.K., J.N., G.E. and M.S. conceived and supervised the project. M.S., A.K. and J.N. wrote the article with contributions from all authors.

